# Visual-like 2D Geometric Template Diffusion for Boosting Single-Sequence Protein Structure Prediction

**DOI:** 10.1101/2025.07.03.662909

**Authors:** Xudong Wang, Tong Zhang, Zhen Cui, Xu Guo, Fuyun Wang, Yuanzhi Wang, Xing Cai, Wenming Zheng

## Abstract

Single-sequence protein structure prediction has drawn increasing attention due to the high computational costs associated with obtaining homologous information. Here, we propose *a visual-like 2D geometric template*^∗^ *diffusion method, named TDFold, to generate high-quality pairwise geometries (including pairwise distances and orientations) for achieving accurate and highly efficient single-sequence 3D structure prediction for proteins.* Given a protein sequence, TDFold initially generates high-quality inter-residue geometries from a probabilistic diffusion perspective. Since inter-residue geometries can be encoded as multi-channel feature matrices, analogous to image feature maps, we construct an image-level 2D geometric template diffusion module by adapting the stable diffusion (SD) model from text-vision generation to sequence-geometry diffusion for proteins. Subsequently, a lightweight sequence-geometry collaborative learning (SCL) network is constructed to facilitate accurate and efficient protein structure prediction. As a result, TDFold possesses three highlights: (i) *better single-sequence prediction performance*: TDFold greatly outperforms existing protein language models (PLMs, e.g. ESMFold and OmegaFold) and homology-based methods (e.g. AlphaFold2, AlphaFold3 and RoseTTAFold) on homology-insufficient datasets such as Orphan and Orphan25, while also achieving promising results on the popular CASP14, CASP15 and CASP16 benchmarks; (ii) *low resource consumption*: By utilizing the lightweight SCL architecture, the GPU memory consumption of TDFold is generally lower than that of popular methods such as AlphaFold2 and ESMFold; (iii) *higher efficiency in training and inference*: TDFold can be trained within a week using a single NVIDIA 4090 GPU. Furthermore, the inference time of TDFold is significantly shorter (about 10x to 100x) than that of existing methods (ESMFold, AlphaFold2 and AlphaFold3) for long protein sequences. This work demonstrates the effectiveness of leveraging powerful vision diffusion models to enhance protein 2D geometric template generation, thereby establishing a new paradigm for single-sequence protein structure prediction. It also accelerates protein-related research, particularly for resource-limited universities and academic institutions. The code has been released to speed up biological research.

## 1 Introduction

In recent years, significant advancements have been achieved in AI-based protein structure prediction ^[1–9]^. Notably, AlphaFold2 ^[1]^, AlphaFold3 ^[10]^ and RoseTTAFold ^[2]^ have demonstrated exceptional performance on the CASP14 dataset ^[11]^, marking a new milestone in the field. However, these deep models heavily rely on homologous information, including multiple sequence alignments (MSAs) ^[12]^ and 3D structural templates ^[13]^, which are typically searched in biological databases such as UniRef ^[14]^ and PDB ^[15]^. This reliance results in low accuracy of homology-based methods for proteins with few or limited homologous information. As illustrated in Fig. 1, 3D structural templates are essential for defining the relative positions of residues, and the exclusion of 3D structural template information leads to significant performance degradation for AlphaFold2 and RoseTTAFold. Additionally, deep learning methods based on protein language models (PLMs) ^[16–20]^, such as ESMFold ^[16]^, trRosettaX-single ^[17]^, RGN2 ^[18]^, and OmegaFold ^[19]^, have recently emerged. These approaches leverage only the textual context of amino acid sequences, eliminating the need for homologous information and accelerating the protein feature extraction process.

**Fig. 1.**
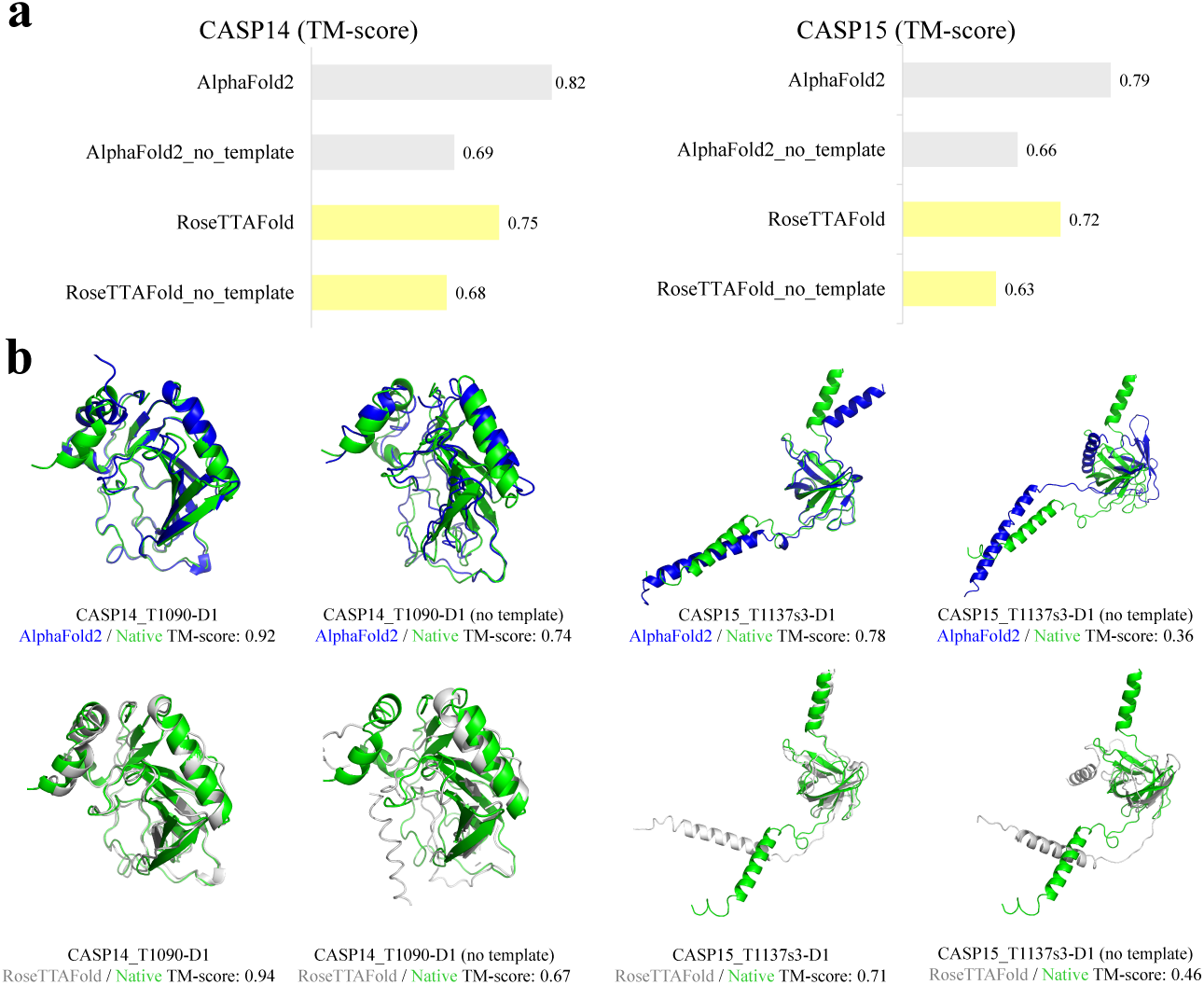
The importance of 3D structural template information to AlphaFold2’s and RoseTTAFold’s protein structure prediction performance on the CASP14 and CASP15 datasets. **a,** TM-score difference between two popular models (AlphaFold2, RoseTTAFold) and their “no template” versions on CASP14 and CASP15 datasets. **b,** The visualization results of 3D structure comparison between the original and “no template” versions. The AlphaFold2’s predicted structures, the RoseTTAFold’s predicted structures and the native structures are painted in blue, gray, and green, respectively.

Despite the significant advancements achieved by homology-based and PLM-based methods, two critical challenges persist in enhancing the research and application of these protein structure prediction models. First, current homology-based methods rely heavily on the availability of homologous data. Consequently, prediction accuracy degrades markedly for targets with few or no homologs—such as orphan proteins or rapidly evolving viral proteins. Moreover, the requisite generation of MSA and 3D structural template identification entails computationally intensive database searches, creating a bottleneck in both speed and scalability. Second, existing PLM-based methods adopt a large-scale architecture derived from language models, which consists of deeply stacked transformer blocks ^[21]^. Notably, the time complexity of the triangular attention mechanism used in ESMFold’s evoformer is *O*(*n*^3^), where *n* denotes the sequence length. This quadratic scaling relationship makes such methods resource-intensive and computationally expensive, leading to substantial memory requirements and computational overhead, particularly for long-sequence proteins.

In this study, we present TDFold, a novel framework that introduces large visual generative models to protein structure prediction, establishing a new technological path for single-sequence-based structure prediction. Given an amino acid sequence, TDFold infers its 3D structure through an end-to-end network architecture consisting of two distinct stages: 2D geometric template diffusion and sequence-geometry collaborative learning (SCL). Specifically, the 2D geometric template diffusion stage aims to derive high-quality interresidue geometries (e.g., matrices of inter-residue distances and orientations) to enhance single-sequence structure prediction. Recognizing that inter-residue geometries can be encoded as multi-channel, image-like feature matrices ^[22]^, we adapt the Stable Diffusion (SD) framework ^[23]^ to develop a diffusion-based method for generating inter-residue geometries directly from sequence. Concretely, a protein 2D geometric template diffusion module is proposed by constructing two Low-Rank Adaptation (LoRA) ^[24]^ branches attached to the text and image (i.e., UNet ^[25]^) encoders in the SD model. Leveraging the SD model’s powerful ability to model collaborations between textual sequences and multi-channel matrices, our 2D geometric template diffusion module enables reliable inter-residue geometries generation (i.e., “images” describing inter-residue distances and orientations) by treating given amino acid sequences as “text prompts”. In the subsequent SCL stage, a lightweight graph network ^[26]^ is constructed to predict protein structures by fusing a residue-level branch and another atom-level one. Specifically, the residue-level learning branch models sequence and inter-residue collaborations, while the atom-level graph learning branch captures the influence of side-chain atoms on backbone conformations within an amino acid sequence. These two branches work collaboratively to derive the final 3D coordinates of proteins. Compared to the existing methods, TDFold offers three key advantages: (i) Superior single-sequence prediction performance: By leveraging the powerful text-to-image generation capability of the SD model and using real inter-residue geometries as supervised information, TDFold generates high-quality inter-residue geometries as intermediate features. This eliminates dependence on homology data and significantly improves the performance of single-sequence protein structure prediction. (ii) Low resource consumption: By adopting the lightweight SCL architecture, TDFold’s GPU memory consumption is generally lower than that of popular protein structure prediction methods such as AlphaFold2 and ESM-Fold. (iii) Higher efficiency in training and inference: TDFold can be trained within one week using a single NVIDIA 4090 GPU, including fine-tuning the SD model and training the SCL network from scratch. Furthermore, the inference time of the SD model is significantly shorter (approximately 10x to 100x) than that of PLM-based methods, particularly for long-sequence proteins.

We report the performance on the homology-insufficient Orphan ^[18]^ and Orphan25 ^[17]^ datasets, as well as the popular CASP14, CASP15 and CASP16 datasets (Critical Assessment of Techniques for Protein Structure Prediction ^[11, 27, 28]^). The experimental results demonstrate that our TDFold achieves state-of-the-art performance on the homology-insufficient datasets Orphan and Orphan25, as well as competitive performance (measured by TM-score ^[29]^, GDT TS ^[30]^ and pLDDT ^[1, 31]^) compared to ESMFold and OmegaFold on CASP14, CASP15 and CASP16. Furthermore, we provide an analysis of the computational time and GPU memory requirements for AlphaFold2, AlphaFold3, RoseTTAFold, ESMFold, and TDFold in protein structure prediction. TDFold requires approximately 10 seconds to predict the structure of a protein containing 500 residues, while ESMFold requires about 100 seconds, AlphaFold3 needs about 240 seconds, AlphaFold2 and RoseTTAFold require nearly 1000 seconds. In terms of GPU memory usage, TDFold occupies about 7 GB, whereas AlphaFold2, RoseTTAFold, and ESM-Fold require 12 GB, 16 GB, and 20 GB, respectively (due to the lack of open training weights for AlphaFold3, we used the AlphaFold3’s server for testing and were unable to obtain its GPU memory usage data). Additionally, we visualize multiple cases of TDFold’s predicted 3D structures and compare the generated inter-residue distances with those derived from homologous 3D structural templates searched in biological databases. These experiments and model analyses collectively demonstrate that TDFold achieves the best protein structure prediction performance in a single-sequence-based mode while requiring the least memory and time for model inference.

## 2 Results

We firstly summarize the framework of the proposed TDFold. Then, we conduct comprehensive experiments on multiple popular protein datasets, including Orphan, Orphan25, CASP14, CASP15 and CASP16. Finally, we analyze the effectiveness of each module in TDFold in the ablation study.

### Approach Summary

The overall framework of the proposed TDFold is depicted in Fig. 2. This model employs a two-stage architecture for three-dimensional structure prediction. Initially, it generates inter-residue geometric information using a 2D geometric template diffusion module, followed by a lightweight sequence-geometry collaborative learning (SCL) network to predict the 3D structure. Specifically, for a given amino acid sequence, the 2D geometric template diffusion module denoises Gaussian distributed noises to generate the inter-residue geometry images (i.e. distance and orientation matrices) while using the protein sequence as the text prompt. To address the significant datatype difference (e.g. continuous geometries vs. discrete images) and semantic gap (e.g. 2D geometric template information vs. general images), we firstly discretize the value of the inter-residue geometric matrices and map them to the pixel value range (0-255) of the RGB image ^[32]^. Then, we apply the Low-Rank Adaptation (LoRA) ^[24]^ fine-tuning technique to the stable diffusion (SD) model ^[23]^. Specifically, the original training parameters of the SD model are frozen, and only the parameters of LoRA are trained in the fine-tuning process. For text encoder ^[33]^, we take a text LoRA to align sequence textual features with inter-residue geometric image features, mapping them into a shared latent space. For the UNet ^[25]^, we utilize four UNet LoRAs focusing on learning the matrices of the distance of C*_β_*-C*_β_*, two dihedrals *ω* (C*_α_*-C*_β_*-C*_β_*-C*_α_*), *θ* (N-C*_α_*-C*_β_*-C*_β_*) and a planar angle *ϕ*: C*_α_*-C*_β_*-C*_β_*, to enhance the model’s capacity to accurately learn the inter-residue geometries. With the help of LoRA fine-tuning technique, we transfer the SD model with powerful image generation capability to the inter-residue geometries generation task. Through the denoising UNet, the SD model is able to capture the complex distribution of inter-residue geometric images and generate reliable samples as 2D geometric templates in the prediction process.

**Fig. 2.**
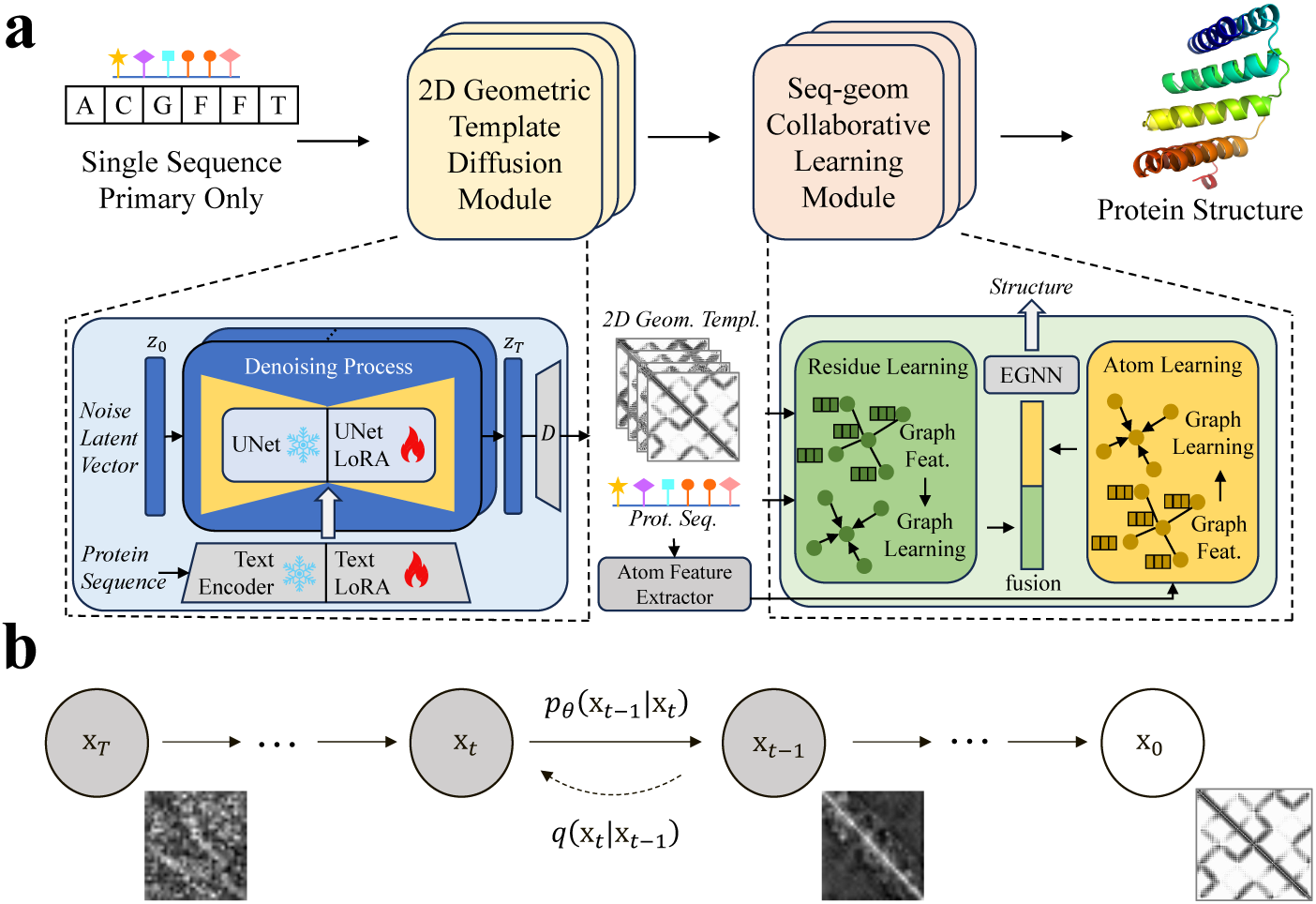
The architecture and 2D geometric template diffusion process of TDFold. **a,** The architecture of TDFold for protein structure prediction. TDFold consists of two modules, the 2D geometric template diffusion module and the sequence-geometry collaborative learning (SCL) module. For one given amino acid sequence, the 2D geometric template diffusion module generates inter-residue geometries guided by the sequence. Next, the sequence and generated inter-residue geometries are sent into the SCL module. The SCL learning module has a two-branch architecture to learn residue-level and atom-level features, respectively. Finally, the residue and atom features are fused and sent into the SE(3) equivariant graph neural network (EGNN) to predict the 3D structure. **b,** The forward and reverse process of the 2D geometric template diffusion in TDFold. The forward process *q* of is a Markov chain that continuously adds Gaussian noises until the original inter-residue geometric image signal is covered by the noise signal. The reverse process *p_θ_* learns to gradually denoise a normally distributed variable and restore the inter-residue geometric image.

Next, we establish the SCL network comprising four key components: residue-level learning, atomic-level learning, residue-atom graph feature fusion, and EGNN-based protein all-atom coordinate prediction. The residue-level branch learns sequence-geometry interactions through the residue features and their pairwise geometric relationships. These outputs are structured as a graph, where residues form nodes and their inter-residue relationships form edges. A graph neural network (GNN) ^[34]^ then learns the residue representations by propagating information across these edges. Concurrently, an atom-level branch explicitly incorporates side-chain effects by constructing a graph of atoms (nodes) and bonds (edges). An atomic GNN ^[35]^ processes this graph to learn representations that integrate both backbone and side-chain atomic features. The two branches are fused via a variational learning framework, which injects side-chain awareness into the backbone representation, enabling the adaptive refinement of residue-level features based on finegrained atomic variations. The resulting integrated representation is finally passed to an SE(3)-equivariant graph neural network (EGNN) ^[36]^ to predict the full atomic 3D structure.

### Predicting structures of orphan proteins

To evaluate our TDFold to orphan proteins, we test the performance on Orphan and Orphan25 datasets which typically contain limited or no homologous information. Also, we compare the results with the three state-of-the-art methods named AlphaFold2 ^[1]^, AlphaFold3 ^[10]^ and RoseTTAFold ^[2]^, as well as four protein language model (PLM) based methods: ESMFold, OmegaFold, RGN2 and trRosettaX-single ^[16–19]^. Specifically, only single sequence mode is tested for PLM-based methods and our TDFold, while both modes of using single sequence and full homology (including MSA and 3D structural template) are tested for AlphaFold2 and RoseTTAFold (only full homology mode for AlphaFold3). For performance measurement, the widely adopted TM-score ^[29]^, GDT TS ^[30]^ and pLDDT ^[1, 31]^ are used as the metric for protein structure prediction.

As shown in Fig. 3a, our TDFold outperforms all these comparison methods on both Orphan and Orphan25 datasets^1^. Specifically, for homology-based methods (AlphaFold2, AlphaFold3, RoseTTAFold), our TDFold outperforms them even though they use the full MSA and 3D structural templates (full MT). This is because the homology-based methods highly rely on homologous information as input, while orphan proteins have rather limited homology (MSA and 3D structural template). Among the PLM-based methods (ESM-Fold, OmegaFold, RGN2, and trRosettaX-single), the structure prediction performance of ESMFold is better than the other three. Compared with them, TDFold achieves the best performances and outperforms ESMFold with the TM-score gains of 0.04 on Orphan and 0.07 on Orphan25. This may be attributed to the fact that TDFold well establishes a distribution mapping between one single protein sequence and the corresponding inter-residue geometries, which effectively boosts the protein structure prediction. Concretely, on the Orphan dataset, the average TM-score of TDFold is 0.46, while that of ESMFold is 0.42, OmegaFold is 0.39 (with the p-value of 0.0227 compared to TDFold), AlphaFold2 (full MT) is 0.37, AlphaFold3 (full MT) is 0.41, and RoseTTAFold (full MT) is 0.35. And for the Orphan25 dataset, the TMscore values are 0.61 of TDFold, 0.54 of ESMFold, 0.52 of OmegaFold (with the p-value of 0.0256 compared to TDFold), 0.44 of AlphaFold2 (full MT), 0.57 of AlphaFold3 (full MT), and 0.40 of RoseTTAFold (full MT).

**Fig. 3.**
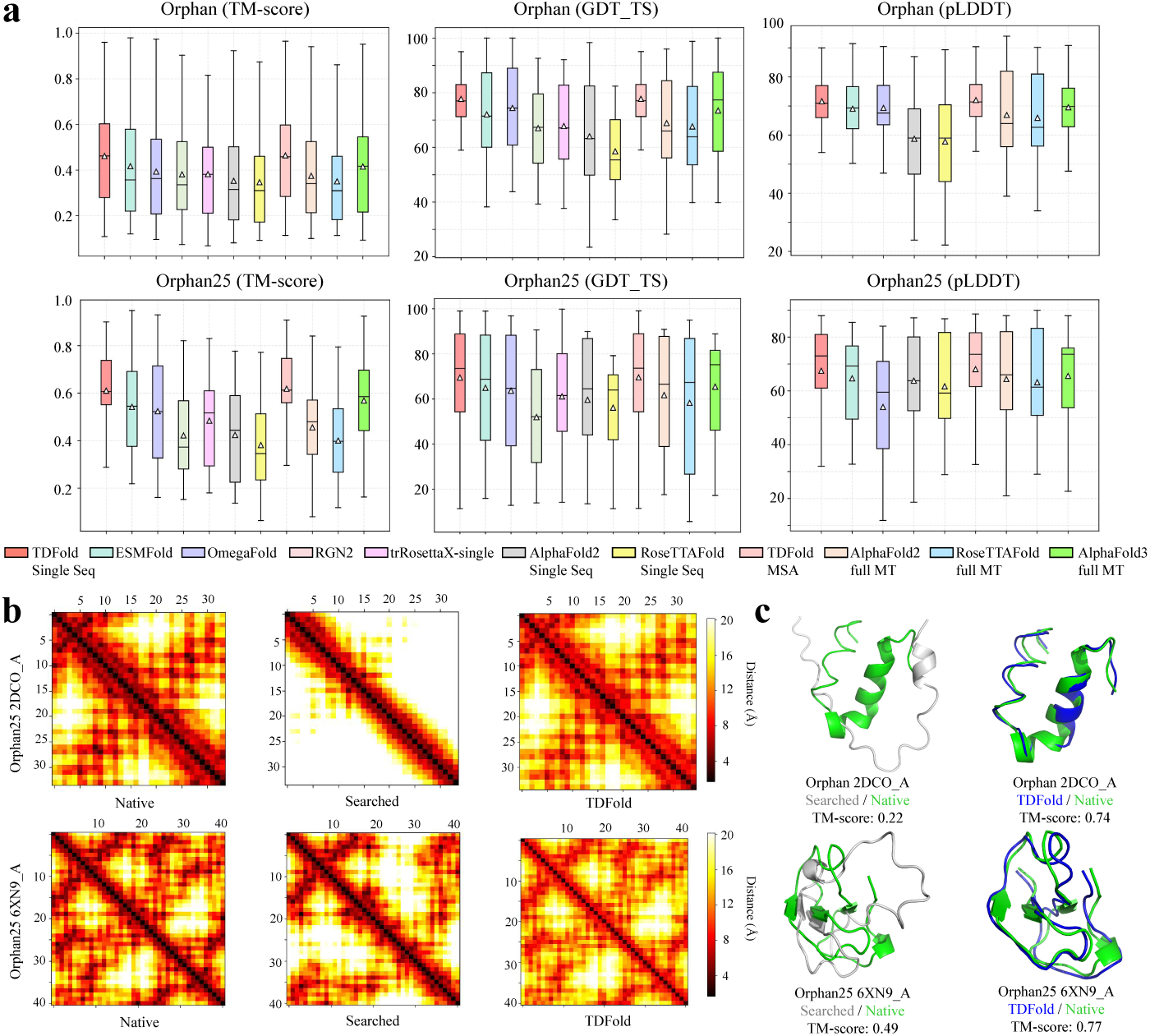
The performance comparison on the Orphan and Orphan25 datasets and the visualization examples. **a,** The performances (TM-score, GDT TS and pLDDT) on the Orphan and Orphan25 datasets of TDFold (Single Seq), ESMFold, OmegaFold, RGN2, trRosettaX-single, AlphaFold2 (Single Seq), RoseTTAFold (Single Seq), TDFold (MSA), AlphaFold2 (full MSA and 3D structural template, full MT), RoseTTAFold (full MT), and AlphaFold3 (full MT) (since RGN2 and trRosettaX-single do not support the calculation of pLDDT, the pLDDT results shown in the figure exclude theirs.). For each box in the figure, the center line, bottom line, and top line represent the median, first quartile, and third quartile, respectively. The horizontal lines along the top and bottom edges represent the maximum and minimum observations. Besides, the white triangle represents the average value. **b,** The inter-residue distance matrices for 2DCO A (Orphan) and 6XN9 A (Orphan25) computed from the native 3D structures, the searched 3D template structures, and those generated by TDFold. Each data point in the distance image corresponds to a pair of residues. The distances are represented by a color gradient and the darker shades indicate closer proximity. In addition, the sequence similarity between native protein and template protein are 0.25 for 2DCO A and 0.29 6XN9 A, respectively. **c,** The TDFold’s prediction (in blue), the searched 3D template’s structure (in gray) and the native structure (in green) of 2DCO A of Orphan and 6XN9 A of Orphan25.

For the another metric GDT TS, on the Orphan dataset, the mean value of TDFold is 77.50, while that of ESMFold is 72.08, OmegaFold is 74.44, AlphaFold2 (full MT) is 68.91, AlphaFold3 (full MT) is 73.49, and RoseTTAFold (full MT) is 67.61. For the Orphan25 dataset, the GDT TS values are 68.37 of TDFold, 64.93 of ESMFold, 63.65 of OmegaFold, 61.70 of AlphaFold2 (full MT), 65.46 of AlphaFold3 (full MT), and 58.25 of RoseTTAFold (full MT). Moreover, we also employ the pLDDT metric to assess the confidence of the model’s predicted structures. The results of TDFold, ESMFold, OmegaFold, AlphaFold2 (full MT), AlphaFold3 (full MT) and RoseTTAFold (full MT) on Orphan dataset are 71.85, 69.52, 68.75, 67.23, 69.55, and 65.96, respectively. On the Orphan25 dataset, the pLDDT values are 67.48 of TDFold, 64.65 of ESMFold, 54.25 of OmegaFold, 64.87 of AlphaFold2 (full MT), 65.57 of AlphaFold3 (full MT), and 63.25 of RoseTTAFold (full MT). According to the results, the pLDDT confidence scores of TDFold provide a good indication of the agreement with native structures.

Additionally, we provide the visualizations of the inter-residue distance matrices (2DCO A from Orphan and 6XN9 A from Orphan25) of the searched 3D structural template and TDFold in Fig. 3b. Meanwhile, the predicted structures of TDFold that use the searched and TDFold-generated geometries as SCL’s inputs are depicted in Fig. 3c. For 2DCO A and 6XN9 A, only a few homologous 3D structural templates can be searched, and they present obvious differences from the native structure according to Fig. 3b. In contrast, the inter-residue distance images generated by TDFold are both more similar with the native ones. Correspondingly, using the generated inter-residue geometries for prediction obtains much better performances than using the searched 3D structural templates. Moreover, the Table 1 displays the results (measured by Kullback-Leibler (KL) divergence ^[37]^, ranging from **0** ∼ +∞) between the inter-residue geometries generated by TDFold and native labels, as well as the KL-divergence results of geometries predicted by trRosetta^[8]^. From the experimental results, it can be seen that the TDFold outperforms trRosetta on Orphan and Orphan25 datasets. As shown in Fig. 3c, the TM-scores of using generated inter-residue geometries are 0.74 (2DCO A) and 0.77 (6XN9 A), which are much higher than 0.22 and 0.49 of using the searched ones.

**Table 1.**
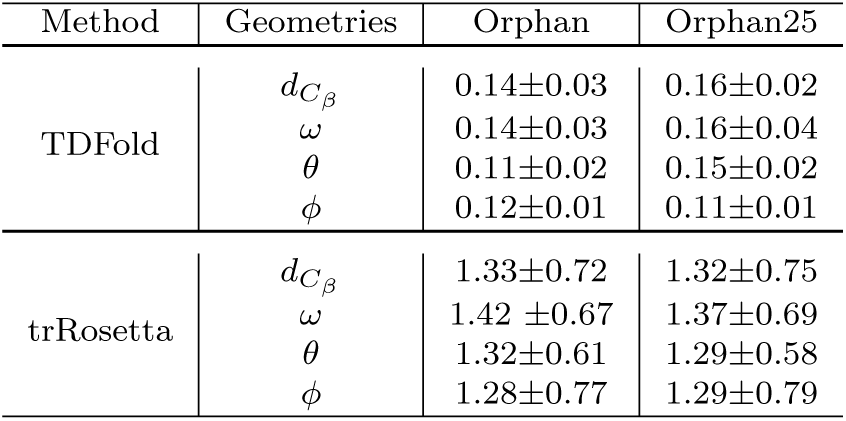
The KL-divergence of inter-residue geometries generated by TDFold and predicted by trRosetta on Orphan and Orphan25 datasets.

| Method | Geometries | Orphan | Orphan25 |
| --- | --- | --- | --- |
| TDFold | $d_{C\beta}$ | 0.14±0.03 | 0.16±0.02 |
| | $\omega$ | 0.14±0.03 | 0.16±0.04 |
| | $\theta$ | 0.11±0.02 | 0.15±0.02 |
| | $\phi$ | 0.12±0.01 | 0.11±0.01 |
| trRosetta | $d_{C\beta}$ | 1.33±0.72 | 1.32±0.75 |
| | $\omega$ | 1.42 ±0.67 | 1.37±0.69 |
| | $\theta$ | 1.32±0.61 | 1.29±0.58 |
| | $\phi$ | 1.28±0.77 | 1.29±0.79 |

### Predicting protein structures in CASP14, CASP15 and CASP16

To evaluate the performance of the TDFold for general protein structure prediction, we compare it with three homology-based methods named AlphaFold2 ^[1]^, AlphaFold3 ^[10]^ and RoseTTAFold ^[2]^, and two PLM-based method called ESMFold ^[16]^ and OmegaFold ^[19]^ on the popular CASP14, CASP15 and CASP16 datasets. The experimental results are shown in Fig. 4a. Overall, our TDFold achieves high prediction performances (average TM-score, GDT TS, and pLDDT) on the CASP14, CASP15, and CASP16 datasets^2^. For AlphaFold2 and RoseTTAFold, they obtain the TM-scores of 0.80, 0.75 on CASP14, 0.79, 0.68 on CASP15, and 0.78, 0.76 on CASP16 with full MSAs and 3D structural templates as inputs. However, when using single sequences as inputs, their performances degrade to the TM-scores of 0.46, 0.43 on CASP14, 0.51, 0.44 on CASP15, and 0.50, 0.46 on CASP16, respectively. For PLM-based methods like ESMFold and OmegaFold, our TDFold is better than ESMFold by obtaining 0.02 (0.73 of TDFold vs. 0.71 of ESMFold) TM-score performance gain on CASP14, 0.01 (0.7 of TDFold vs. 0.69 of ESMFold) gain on CASP15, and 0.02 (0.77 of TDFold vs. 0.75 of ESMFold) gain on CASP16. In addition, as shown in Fig. 4a, our method significantly outperforms OmegaFold by achieving 0.07 TM-score gain (0.7 of TDFold vs. 0.63 of OmegaFold) on the CASP15 dataset (p-value 0.0268), 0.08 TM-score gain (0.77 of TDFold vs. 0.69 of OmegaFold) on the CASP16 dataset (p-value 0.0246), while obtaining a performance that is 0.03 lower (0.73 of TDFold vs. 0.76 of OmegaFold) on the CASP14 dataset (p-value 0.0275). It should be noticed that the training datasets of OmegaFold contain the protein sequences UniRef50 dataset (dated at 2021/04), and the structures from the Protein Databank PDB deposited before 2021, while the protein structures of CASP14 were released in the Protein Databank PDB from 2020/07. In contrast, the released training dataset dates of AlphaFold2, RoseTTAFold, ESMFold, and our TDFold are 2020/05.

**Fig. 4.**
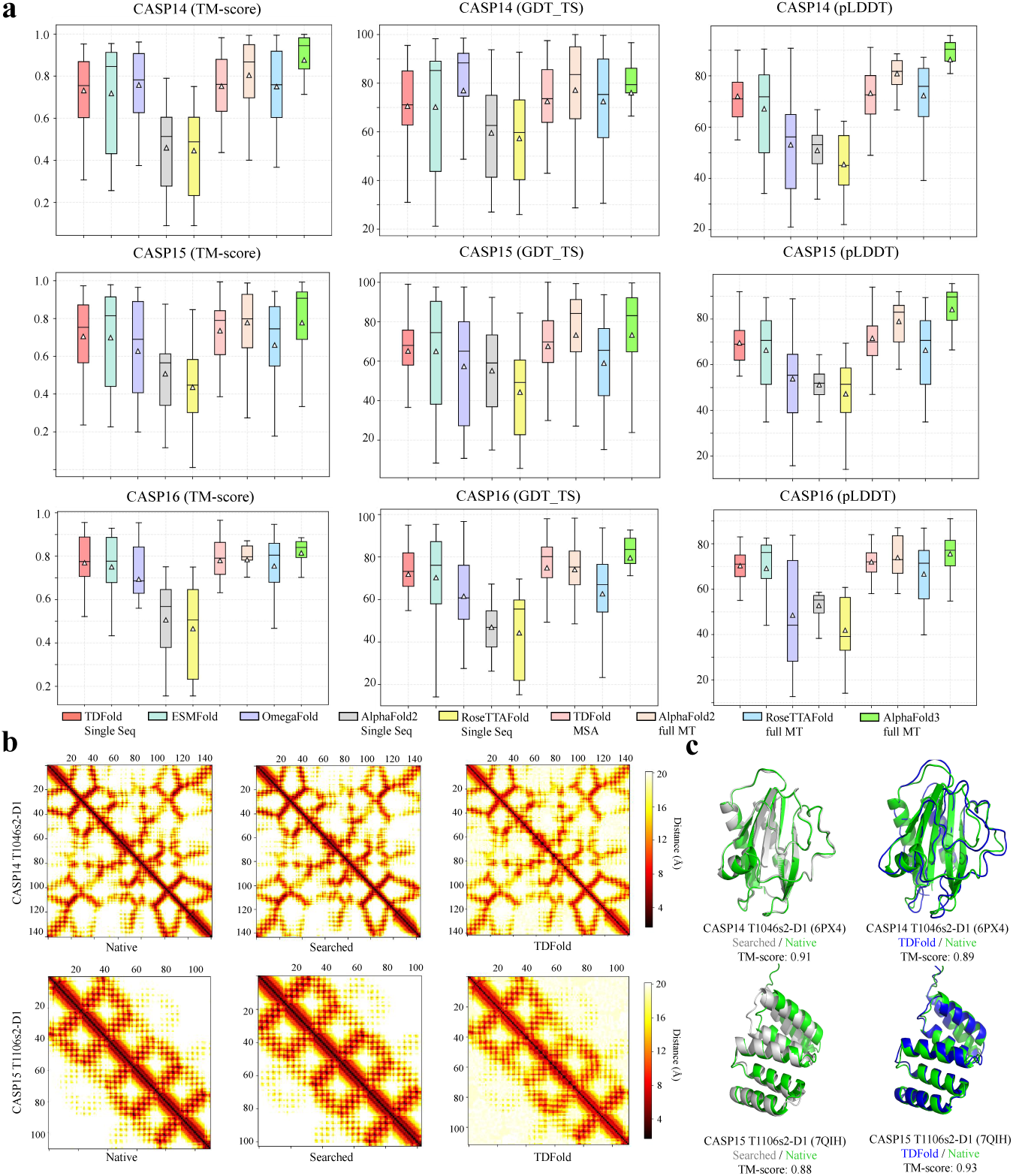
The performance comparison on the CASP14, CASP15 and CASP16 datasets and the visualization examples. **a,** The TM-score, GDT TS, and pLDDT values of TDFold (Single Seq), ESMFold, OmegaFold, AlphaFold2 (Single Seq), RoseTTAFold (Single Seq), TDFold (MSA), AlphaFold2 (full MT), RoseTTAFold (full MT), and AlphaFold3 (full MT) on the CASP14, CASP15 and CASP16 datasets. For each box in the figure, the center line, bottom line, and top line represent the median, first quartile, and third quartile, respectively. The horizontal lines along the top and bottom edges represent the maximum and minimum observations. Besides, the white triangle represents the average value. **b,** The inter-residue distance matrices for T1046s2-D1 (CASP14, PDB code 6PX4) and T1106s2-D1 (CASP15, PDB code 7QIH) computed from the native 3D structures, the searched 3D template structures, and those generated by TDFold. Each data point in the distance image corresponds to a pair of residues. The distances are represented by a color gradient and the darker shades indicate closer proximity. In addition, the sequence similarity between native protein and template protein are 0.85 for T1046s2-D1 and 0.71 for T1106s2-D1, respectively. **c,** The TDFold’s prediction (in blue), the searched 3D template’s structure (in gray) and the native structure (in green) of T1046s2-D1 and T1106s2-D1.

To further comprehensively measure the performance, we also employ the GDT TS and pLDDT metrics for evaluation. The GDT TS results of TDFold, ESMFold, OmegaFold, AlphaFold2 (full MT), AlphaFold3 (full MT), and RoseTTAFold (full MT) on the CASP14 dataset are 70.86, 70.21, 75.35, 76.15, 77.08 and 72.07, respectively. On the CASP15 dataset, the GDT TS values are 63.52 of TDFold, 62.99 of ESMFold, 57.37 of OmegaFold, 73.24 of AlphaFold2 (full MT), 73.26 of AlphaFold3 (full MT), and 60.35 of RoseTTAFold (full MT). And for the CASP16 dataset, the GDT TS scores are 71.91 of TDFold, 70.33 of ESMFold, 61.55 of OmegaFold, 74.05 of AlphaFold2 (full MT), 79.59 of AlphaFold3 (full MT), and 62.68 of RoseTTAFold (full MT). Moreover, we also compare several methods with our TDFold using the pLDDT metric to assess the models’ prediction confidence. On the CASP14 dataset, the mean value of TDFold is 72.06, while ESMFold is 67.14, OmegaFold is 53.25, AlphaFold2 (full MT) is 80.5, AlphaFold3 (full MT) is 81.35, and RoseTTAFold (full MT) is 72.3. For the CASP15 dataset, the pLDDT values are 69.6 of TDFold, 66.38 of ESMFold, 53.83 of OmegaFold, 79.25 of AlphaFold2 (full MT), 84.21 of AlphaFold3 (full MT), and 66.35 of RoseTTAFold (full MT). And on the CASP16 dataset, the pLDDT values are 70.33 of TDFold, 69.11 of ESMFold, 48.52 of OmegaFold, 73.86 of AlphaFold2 (full MT), 75.53 of AlphaFold3 (full MT), and 66.64 of RoseTTAFold (full MT). The approximate performances between the GDT TS and pLDDT metrics further verify the good agreement with native structures of TDFold’s predictions.

We visualize several inter-residue distance matrices for the proteins T1046s2-D1 (PDB code 6PX4) from CASP14 and T1106s2-D1 (PDB code 7QIH) from CASP15 in Fig. 4b. For T1046s2-D1 and T1106s2-D1, both the generated inter-residue distance images of TDFold and searched 3D structural templates are closely aligned with the native ones. Accordingly, high TM-scores are obtained by using the searched 3D structural templates (0.91 for T1046s2-D1 and 0.88 for T1106s2-D1) and generated ones (0.89 for T1046s2-D1 and 0.93 for T1106s2-D1). In addition, the Table 2 displays the results (measured by KL-divergence) between the inter-residue geometries generated by TDFold and native labels, as well as the KL-divergence results of geometries predicted by trRosetta. From the experimental results, it can be seen that the TDFold outperforms trRosetta on all CASP datasets. From the experimental results, it can be seen that our TDFold is able to generate high-quality inter-residue geometries comparable with those computed from the highly matched 3D structural templates, which verifies the effectiveness of the designed paradigm of transferring the visual SD model to the protein 2D geometric template generation.

**Table 2.** The KL-divergence of inter-residue geometries generated by TDFold and predicted by trRosetta on CASP14, CASP15 and CASP16 datasets.

| Method | Geometries | CASP14 | CASP15 | CASP16 |
| --- | --- | --- | --- | --- |
| TDFold | $d_{C_\beta}$ | 0.12±0.01 | 0.15±0.01 | 0.14±0.02 |
| | $\omega$ | 0.25±0.02 | 0.23±0.02 | 0.22±0.03 |
| | $\theta$ | 0.11±0.02 | 0.13±0.04 | 0.14±0.02 |
| | $\phi$ | 0.18±0.02 | 0.16±0.02 | 0.15±0.01 |
| trRosetta | $d_{C_\beta}$ | 1.16±0.47 | 1.15±0.45 | 1.16±0.43 |
| | $\omega$ | 1.28±0.43 | 1.26±0.42 | 1.23±0.45 |
| | $\theta$ | 1.19±0.44 | 1.18±0.46 | 1.21±0.45 |
| | $\phi$ | 1.18±0.55 | 1.21±0.49 | 1.20±0.52 |

### Promoting the structure prediction of virus-related proteins

Many viruses encode rapidly evolving proteins to evade host immune responses and enhance host-specific adaptation. Notable examples include coronaviruses’ non structural proteins (NSPs) and accessory proteins involved in host immune escape and virus adaptive evolution, typically exhibiting less than 20% sequence homology across strains. These proteins play crucial roles in viral pathogenesis, their low sequence homology presents significant challenges for structural prediction. To evaluate existing methods’ prediction performance on rapidly evolving viruses’ structures, we test them on viral proteins from the CASP14-16 datasets. These included targets with limited homology (*<*20 homologous sequences), such as T1064 – structure of SARS-CoV-2 ORF8 accessory protein. As shown in Table 3, TDFold has superior performance compared to AlphaFold2, AlphaFold3 and ESMFold on these challenging targets. These results highlight TDFold’s enhanced capability for virus structure prediction when homologous sequence information is scarce. TDFold plays a driving role in understanding the mechanisms by which these viruses evade immunity and developing corresponding drugs.

**Table 3.** The TM-score of comparison methods and TDFold on virus-related proteins with a small amount of homologs.

| Protein Type | T1033-D1<br>crAs phage | T1039-D1<br>crAs phage | T1064-D1<br>SARS-CoV-2 | T1082-D1<br>T4 phage | T1099-D1<br>hepatitis virus | T1123-D1<br>astrovirus |
| --- | --- | --- | --- | --- | --- | --- |
| Seq Len | 100 | 161 | 75 | 71 | 178 | 214 |
| MSA Num | 3 | 3 | 14 | 11 | 10 | 9 |
| AlphaFold2 | 0.41 | 0.58 | 0.41 | 0.4 | 0.8 | 0.62 |
| AlphaFold3 | 0.44 | 0.47 | 0.71 | 0.73 | 0.85 | 0.28 |
| ESMFold | 0.33 | 0.28 | 0.44 | 0.38 | 0.47 | 0.31 |
| TDFold | **0.94** | **0.89** | **0.73** | **0.76** | **0.87** | **0.79** |

### Comparison of inference time and GPU memory usage

For largescale structural protein structure prediction tasks, inference time and memory usage are the two major factors determining inference costs. Therefore, we compare the inference computational costs of existing methods to evaluate their applicability for large-scale prediction tasks. Concretely, we provide a detailed comparison of time cost associated with predictions, as illustrated in Fig. 5a. AlphaFold2, AlphaFold3 ^3^ and RoseTTAFold, which rely on the search for homologous protein information, exhibit longer prediction times compared to language models such as ESMFold and our TDFold (e.g. for the protein with sequence length *>* 500, the inference time of AlphaFold2 and RoseTTAFold are both over 1000 seconds, the time of AlphaFold3 is about 240 seconds, the time of ESMFold is about 100 seconds, while the time of TDFold is only about 10 seconds). While ESMFold achieves faster predictions for shorter sequences (*length* ≤ 140), its prediction time escalates with increasing sequence length. This is because ESMFold adopts a triangular attention module with its *O*(*n*^3^) time complexity (where n is the sequence length) in evoformer, which is the primary computational bottleneck governing the model’s prediction time. In contrast, the prediction time of TDFold is primarily determined by the number of denoising steps and independent of the sequence length. And with the help of samplers including DPMsolver ^[38]^, UniPC ^[39]^ and DDIM ^[40]^, the number of denoising steps can be reduced from 1000 to just 25-50, dramatically speeding up the inference process.

**Fig. 5.**
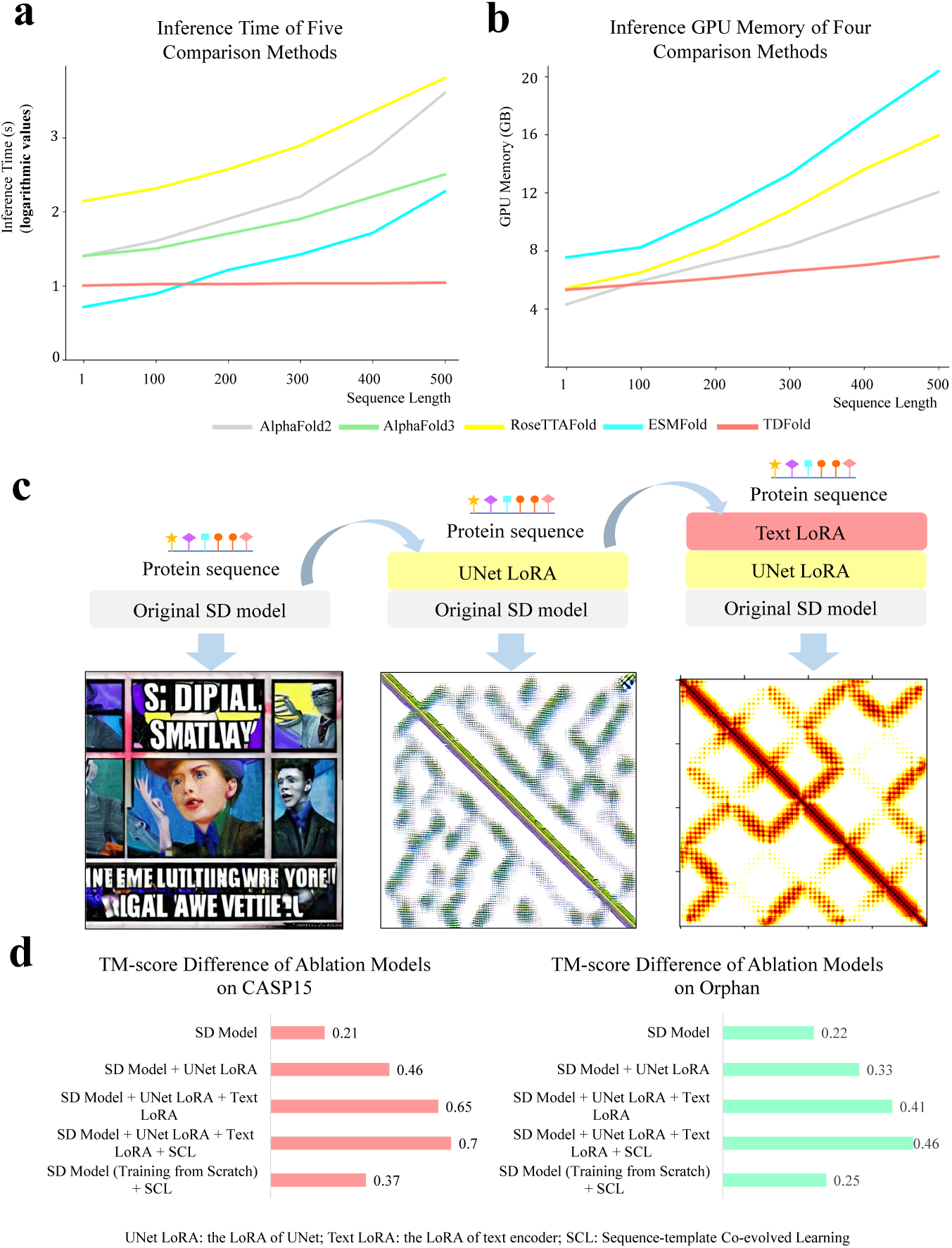
Computation time-memory cost comparisons and model ablation analysis. **a,** Comparison of prediction times among AlphaFold2, AlphaFold3, RoseTTAFold, ESMFold and TDFold across 194 targets spanning our CASP14, CASP15, CASP16, Orphan and Orphan25 protein datasets. The prediction time of TDFold is almost lower than other methods, and only when the sequence length is less than 140, the prediction time is higher than ESMFold. Significantly, the inference time of all other methods increases with the length of the sequence, while the prediction time of TDFold basically remains stable. **b,** The GPU memory used for inference of AlphaFold2, RoseTTAFold, ESMFold, and TDFold. As the length of the amino acid sequence increases, the memory usage of the other three methods increase greatly (AlphaFold2: 4 GB to 12 GB, RoseTTAFold: 5 GB to 16 GB, ESMFold: 8 GB to 20 GB). On the contrary, The memory usage of TDFold has slightly increased from 5 GB to 7 GB. **c,** The differences in the generated inter-residue distance images among the original SD model, the SD model enhanced with the UNet LoRA, and the SD model employing the “Text+UNet” LoRA are presented. Each ablation model is constructed by sequentially adding one or more components to the baseline model. **d,** TM-score difference between the SD model and several ablation models on the CASP15 and Orphan datasets.

Furthermore, the comparison of GPU memory occupation is shown in Fig. 5b. ESMFold exhibits the highest memory usage across all sequences, while AlphaFold2 and RoseTTAFold have moderate GPU memory consumption, and TDFold demonstrates the lowest memory usage on most sequences. Additionally, we analyzed the GPU memory usage during model inference for amino acid sequences of varying lengths. As the length of the amino acid sequence increases, the memory usage of AlphaFold2 and RoseTTAFold increases approximately 3 times (AlphaFold2: 4 GB to 12GB, RoseTTAFold: 5 GB to 16 GB), ESMFold’s memory usage increases by 2.5 times (8 GB to 20 GB), while TDFold’s memory usage only increases by 40% (5 GB to 7 GB). This is because TDFold adopts a lightweight SCL network, reducing the dimensionality and layers of the neural network, thereby reducing the use of GPU memory. Even as sequence length increases, TDFold’s prediction time and GPU memory usage remain almost stable, highlighting the efficiency of TDFold in protein structure prediction.

### Ablation Study

To assess the contribution of each component to TDFold’s performance, we conduct an ablation study illustrated in Fig. 5. Specifically, the effectiveness of UNet LoRA and Text LoRA in promoting inter-residue geometry diffusion is visualized in Fig. 5c, and the performance gains brought by each component on CASP15 and Orphan datasets are shown in Fig. 5d. On the left side of Fig. 5c, when prompted with a protein sequence, the generated image of the original stable diffusion model ^[23]^ presents mainly people and meaningless combinations of characters. This indicates that the model fails to comprehend the semantic context of inter-residue distance matrix and still regards the protein sequence as the natural language describing an image. In contrast, by introducing of the LoRA mechanism into the UNet, the generated image begins to present the diagonal structure, suggesting that the model start to learn the structural layout of inter-residue geometries. However, the sequence embedding and inter-residue distance image embedding are still not aligned, resulting in structural inconsistency in the generated image. Therefore, we add another LoRA for text encoder to finetune the semantics of the protein sequences to make them align with the inter-residue geometric image embeddings. On the right side, after incorporating both the text encoder LoRA and UNet LoRA, the model effectively generates a coherent inter-residue distance image, well capturing the spatial geometric relationships between residues.

As illustrated in Fig.5d, adding each component (i.e. UNet LoRA, Text LoRA, SCL) effectively promote the prediction performance. For UNet LoRA, the considerable performance gains are obtained with 0.25 TM-score on CASP15 and 0.11 TM-score on Orphan. Meanwhile, Text LoRA also effectively improves the performances with 0.19 on CASP15 and 0.08 on Orphan. Furthermore, the SCL module also contributes to the prediction performances where the TM-scores increase approximately both 0.05 for CASP15 and Orphan. Moreover, we also compare the SD model (training-from-scratch + SCL) with the LoRA-finetuned version (UNet LoRA + Text LoRA + SCL) on both the CASP15 and Orphan datasets. The LoRA-finetuned model achieves superior performance, with TM-score improvements of 0.33 on CASP15 (p-value 0.0002) and 0.21 on Orphan (p-value 0.0001). These results demonstrate the importance of fine-tuning – by leveraging the SD model’s pre-trained semantic knowledge from 2 billion image-text pairs, LoRA effectively transfers the knowledge from the text-to-image generation to the inter-residue geometry generation. Overall, in Figure 5, the ablation results verify the effectiveness of each component and the specific importance of UNet LoRA and Text LoRA to boost the accuracy of protein structure prediction. A detailed description of each ablation component is provided in Section 4.

## 3 Discussion

In this study, we introduce TDFold, a protein structure prediction model based on the proposed 2D geometric template diffusion model, and comprehensively evaluate its performance using both quantitative metrics and visualization results. TDFold effectively generates reliable inter-residue geometries from a protein sequence prompt. This approach allows for the utilization of sequence information and the generated inter-residue geometries to predict protein structures, eliminating the time-consuming process of searching for homologous information. Moreover, TDFold demonstrates robustness in handling structure predictions for protein sequences with various levels of homologous information, especially those with a little or even no homologies (e.g., orphan proteins or rapidly evolved viruses related proteins). The comprehensive experimental results on multiple CASP and Orphan datasets also verify their effectiveness in boosting single-sequence protein structure prediction. More importantly, with the help of LoRA fine-tuning technique, TDFold can be deployed on standard personal computers with a single NVIDIA 4090 24GB GPU, facilitating efficient training and inference. The huge advantage of TDFold in inference time also indicates that it is more suitable for large-scale prediction tasks that require high prediction speed.

Overall, TDFold demonstrates superior performance for general proteins or orphan proteins which lack close homologs in databases. Additionally, TDFold is 10–100× faster than AlphaFold2 and requires substantially less GPU memory, making it highly efficient for large-scale protein structure predictions and scenarios with limited computational resources.

## 4 Methods

### Datasets

In this study, the training samples are derived from protein data released in the PDB ^[15]^ prior to May 1, 2020, encompassing a total of 352,409 non-redundant protein domains. We evaluate our proposed model on five datasets: CASP14, CASP15, CASP16, Orphan, and Orphan25. The CASP14 dataset comprises available domain data, with target data correctly split by the domain definition index found in the download area of the CASP14 (Critical Assessment of Techniques for Protein Structure Prediction) website. Ultimately, 32 available domains are included. The CASP15 test set consists of 45 structures obtained from the CASP15 download area. For the latest CASP16 dataset, we use the all released 15 structures. For proteins with insufficient homologous information, we utilize two datasets from previous studies: Orphan ^[18]^ and Orphan25 ^[17]^. The Orphan dataset includes 77 proteins that lack homologs across UniRef30 ^[14]^, PDB70 ^[15]^, and MGnify ^[41]^ simultaneously. Moreover, Orphan25 contains 25 proteins (released from PDB after May 2020) that were searched against the UniRef50 2018 03 database ^[14]^, returning no homologous sequences. The structural similarities (measured by TM-score) between proteins in the training dataset and in the benchmark datasets are all lower than 0.3. We further evaluated the sequence identity using BLAST ^[42]^, revealing that all benchmark datasets share the sequence identity that is less than 30% with the training set.

### The datatype alignment between inter-residue geometries and images for the 2D geometric template diffusion

Due to the inconsistency between the continuous values of inter-residue geometries (distance and orientation) and the discrete image pixel values required by the Stable Diffusion (SD) model ^[23]^, we need to make feature datatype transformation for inter-residue geometric information. Specifically, as shown in Algorithm 1, the inter-residue distance and orientation, including 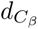 (distance between two “*C_β_*” atoms), two dihedrals: *ω* (C*_α_*-C*_β_*-C*_β_*-C*_α_*), *θ* (N-C*_α_*-C*_β_*-C*_β_*) and a planar angle *ϕ* (C*_α_*-C*_β_*-C*_β_*), define the relative position of any two residues in a protein. For data consistency, we first divide the inter-residue *C_β_* distance range into 36 intervals, i.e., (2.5 Å, 3.0 Å), (3.0 Å, 3.5 Å), · · ·, and (20.0 Å, 20.5 Å). For *ω* and *θ* in orientation, we also divide it into 36 bins from 0 to 360 degrees. But for planar angle *ϕ*, because it ranges from 0 to 180 degrees, it’s divided into 18 parts. After discrete operations, the inter-residue distance and orientation can be represented as N×N (N is the length of protein sequence) discrete matrix with value range from 0 to 36 (0 to 18 for *ϕ*).

#### Algorithm 1 Transformation of inter-residue geometries to images

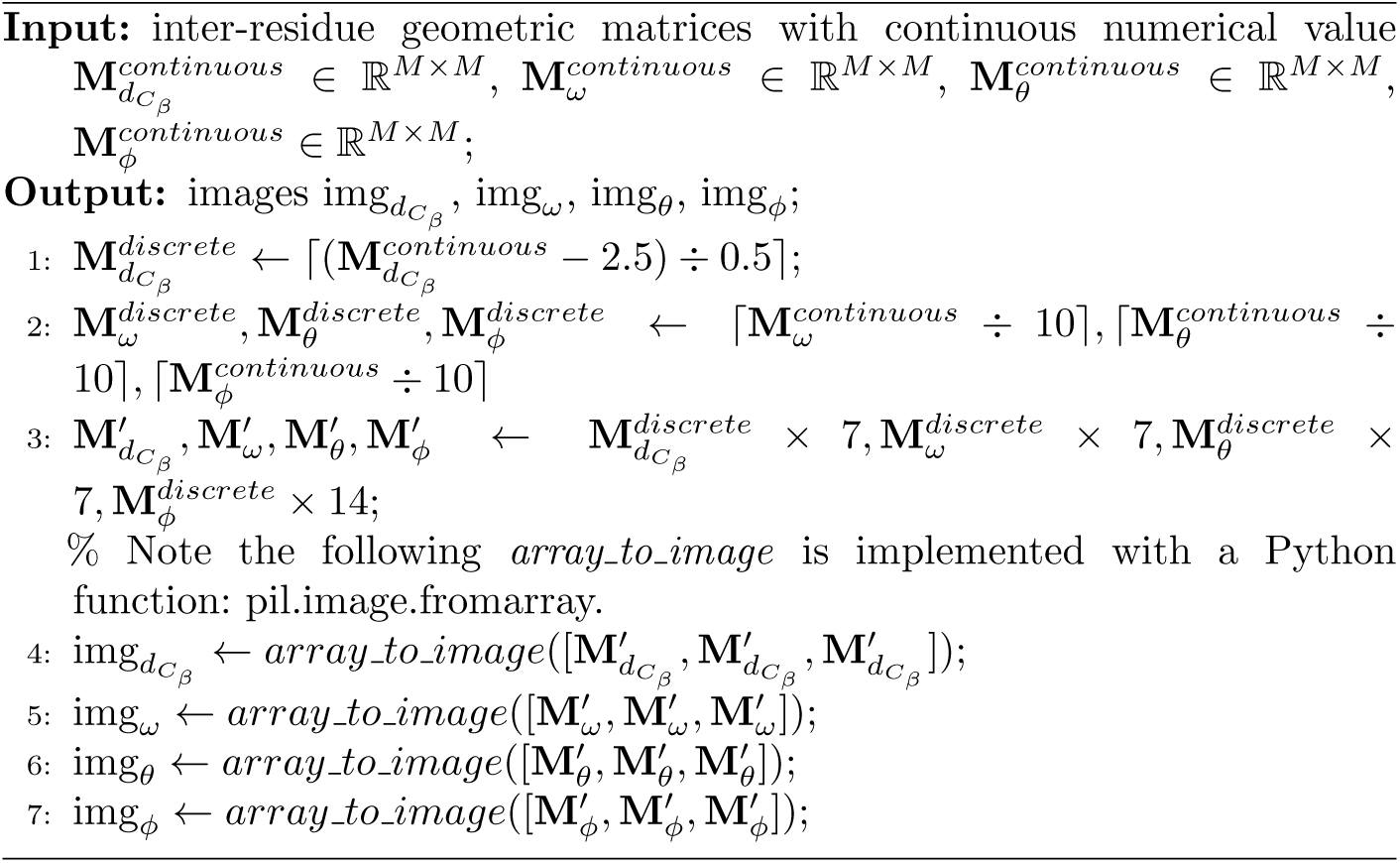

Subsequently, for each component in inter-residue geometries, we independently transform the component into a three-channel image. Concretely, for 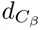, *ω* and *θ*, the matrix values were multiplied by 7 to rescale the range to (0–252), and the resulting matrix is stacked across three channels to form an RGB image. For *ϕ*, matrix values are multiplied by 14 to achieve the same output range (0–252), with all subsequent processing steps identical to those applied to the other parameters. For example, given a N×N distance matrix 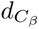, we first multiply each element in the matrix by 7 to expand the numerical range to (0-252) (e. g., 36×7=252), then copy the expanded matrix three times and stack them together as the three channels of the RGB image. In final, the geometric matrix are mapped into the range of RGB image pixel value (0 ∼ 255) and can be processed by SD model.

### The protein 2D geometric template diffusion module for interresidue geometries generation

The protein 2D geometric template diffusion module is constructed through transfer learning based on the SD model. Concretely, the SD model is primarily trained on general images, which means it lacks the specific protein semantic information necessary for accurately representing inter-residue geometric images. To address this gap, we fine-tune the text encoder ^[33]^ and UNet ^[25]^ of the SD model using Low Rank Adaptation (LoRA) ^[24]^ (the specific process is introduced in the next section). This approach enables the model to effectively learn the semantic features associated with protein inter-residue geometries. Finally, the 2D geometric template diffusion module can generate inter-residue geometries with protein sequence prompts as conditions.

Specifically, the 2D geometric template diffusion module is the probabilistic model ^[43]^ designed to learn the inter-residue geometry (distances and orientations) distribution *p*(**x**) (**x** is the inter-residue geometry) by gradually denoising a normally distributed variable, which corresponds to learning the reverse process of a fixed Markov Chain ^[44, 45]^ of length *T*. The probabilistic model can be interpreted as an equally weighted sequence of denoising autoencoders *e_θ_*(**x***_t_, t*); *t* = 1*…T*, which are trained to predict a denoised variant of input **x***_t_*, where **x***_t_* is a noisy version of the input **x**. Furthermore, the 2D geometric template (inter-residue geometries) generation process through the protein sequence prompt *seq* can be regarded as a conditional distributions of the form *p*(**x** | *seq*) and controlled by a conditional denoising autoencoder ^[46]^ *e_θ_*(**x***_t_, t, seq*) with the cross-attention mechanism. To pre-process the protein sequence *seq*, we introduce a protein sequence encoder *τ_θ_* to project the *seq* into an intermediate representation 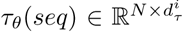. Based on inter-residue geometries and protein sequence condition, we then learn the 2D geometric template diffusion model objective L*_T DM_* via

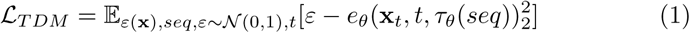

### The hierarchical LoRA model for the 2D geometric template diffusion module

In order to make the SD model applicable to the 2D geometric template diffusion, we introduce the LoRA fine-tuning mechanism to the text encoder and UNet. Specifically, the original training parameters of the SD model are frozen, and only the parameters of LoRA are trained in the fine-tuning process. The integration of LoRA allows the text encoder model to map protein sequences and inter-residue geometries into a shared embedding space. This alignment is crucial for ensuring that the model understands the relationships between protein sequences and their corresponding geometric representations. And we apply LoRA to the UNet, enabling it to learn the distribution of inter-residue geometric data while being guided by protein sequence prompts.

For the text encoder, the incorporation of LoRA facilitates the mapping of protein sequences and inter-residue geometries into a unified embedding space. This integration is essential for enabling the model to effectively capture and interpret complex relationships between protein sequences and their associated geometric matrices. The LoRA model of text encoder can be represented as follows,

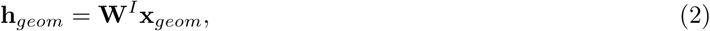

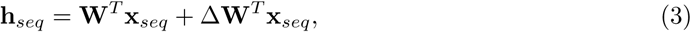

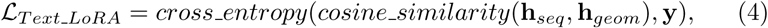

where **W***^T^* is the model weight of the original text encoder, Δ**W***^T^* is the text encoder LoRA weight for learning the collaborations between sequences and inter-residue geometries, and **W***^I^* is the weight of the image encoder of SD model. **x***_seq_* is the sequence embedding and **x***_geom_* is the inter-residue geometry information. Moreover, **h***_seq_* and **h***_geom_* are the hidden states for **x***_seq_* and **x***_geom_*, respectively. Then, we calculate the cross entropy loss between the cosine similarity of sequence features and geometric feature and the ground truth similarity **y**. With the constraint of the loss function, the text features and inter-residue geometries features are gradually mapped into the same latent space.

Next, we apply LoRA to the UNet, enabling it to learn the distribution of inter-residue geometric data while being guided by protein sequence prompts. This process transfers the SD model from general image processing to the specific task of handling inter-residue geometric matrices. By fine-tuning these components, we enhance the model’s ability to capture and generate protein inter-residue geometries effectively. The LoRA model of UNet can be represented as follows,

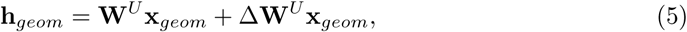

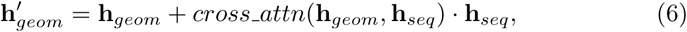

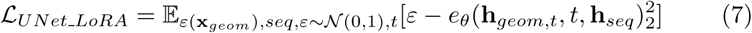

where **W***^U^* is the weight of original UNet model weight, and Δ**W***^U^* is the UNet LoRA weights for predicting the value of inter-residue geometries (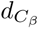*, ω, θ, ϕ*). After semantic alignment of sequence and geometric information through the text encoder LoRA, the sequence information **h***_seq_* is added as a guide to the geometric information generation process by the cross attention. Then, we calculate the MSE loss between the predicted noise *e_θ_*(**h***_geom,t_, t,* **h***_seq_*) and true noise *ε* at each time step *t*. Through optimization of the loss function, the model is trained to estimate the noise profile at each diffusion time step, conditional on the input protein sequence *seq*. This enables a deterministic generative process: starting from isotropic Gaussian noise, the model performs iterative denoising under the specification of the protein sequence *seq* to synthesize its corresponding inter-residue geometric **x***_geom_*. After optimizing the LoRA parameters, the reliable inter-residue geometries can be generated to promote the single-sequence protein structure prediction.

### The long-text encoding capacity for protein sequence prompt embedding

The length of protein sequences varies from tens to thousands. However, the original text encoder used in stable diffusion only supports a maximum of 77 tokens (each token is a amino acid letter)—far fewer than the lengths of typical protein sequences. This token number limitation of the text encoder leads to a rigid constraint on the input protein sequence length.

To overcome the challenge of token numbers in the text encoder, position interpolation is introduced to the positional embedding of the text encoder to enable context window extensions. We leverage the property that position encoding can be extended to non-integer positions by interpolating the encoding at adjacent integer positions ^[47]^. The calculation for obtaining the new extended positional embedding *PE_extend_*(*i*) can be expressed as follows:

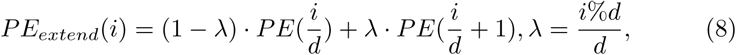

where *PE*(*i*) is the original positional embedding of *i_th_* position, and *d* is the times number of interpolation. *λ* is a ratio between 0 and 1, determining whether the interpolated positional embedding for the *i_th_* position is closer to its preceding or following position.

After the above process, the text length is extended to 385. For the longer sequence, we conduct the non-overlapping segmentation to the sequence and generate embeddings for each segment, and then aggregate the embeddings to generate the final inter-residue geometry.

### The residue-level graph learning

For the sequence-geometry collaborative learning module (SCL), it contains residue-level learning, atomic-level learning, residue-atom graph feature fusion, and EGNN-based protein all-atom coordinate prediction. Specifically, as illustrated in Algorithm 2, the residue-level learning processes the protein sequence and generated inter-residue geometries (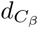*, ω, θ, ϕ*). It comprises three primary learning components: a collaborative learning network, a hybrid convolution neural network (CNN), and a residue graph learning network. Firstly, within the collaborative learning network, interactive learning is conducted between protein sequence and inter-residue geometries. Concretely, the protein sequence is first tokenized and embedded into a feature vector, and pair features are constructed by extracting pairwise distances and relative orientations from the generated inter-residue geometries. Moreover, to ensure that *C_β_* distance and *ω* dihedral matrices satisfy their inherent physical symmetry constraints, taking the original *C_β_* distance matrix 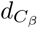 as an example, the symmetric operation 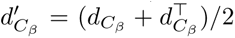 performed to obtain the corresponding symmetric matrix 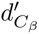.

To model the interplay between these sequence and pair representations, we implement an iterative update mechanism. This mechanism operates bidirectionally: first, an outer product of the sequence feature map is computed and added to the pair representation, thereby updating the pair features with the sequence signals. Second, a cross-attention operation from the sequence to the pair features transfers the structural information encoded in the pairs back to the sequence representation. Furthermore, self-attention layers are applied independently to both the sequence and pair features to enable self-refinement. The output of this process is the initial residue-level features **X***_r_*.

#### Algorithm 2 Residue Branch Graph Learning

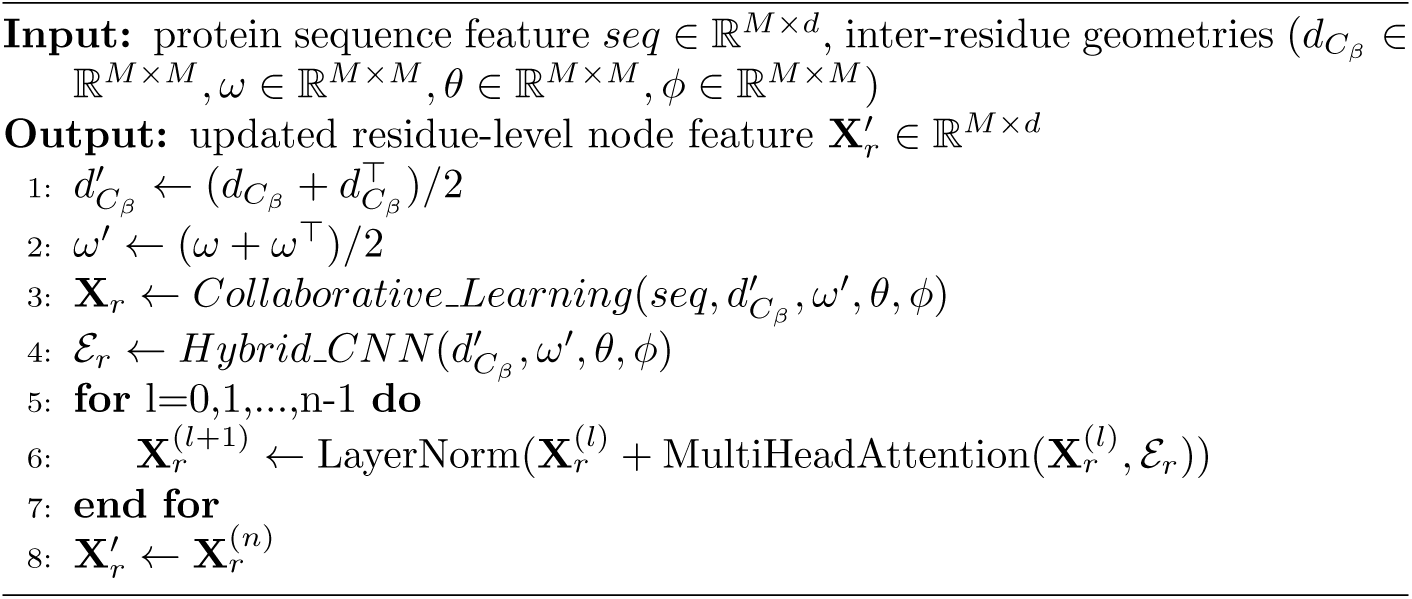

Secondly, the hybrid CNN architecture integrates both asymmetric and symmetric convolution kernels to effectively learn inter-residue geometric features. Asymmetric kernels capture the dihedral angle *θ* (N-C*_α_*-C*_β_*-C*_β_*)and the planar angle *ϕ* (C*_α_*-C*_β_*-C*_β_*), whereas symmetric kernels model the distance 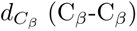 and the dihedral angle *ω* (C*_α_*-C*_β_*-C*_β_*-C*_α_*). This dual-kernel approach produces the residue-level correlation matrix E*_r_* and explicitly preserves the inherent symmetries of the geometric descriptors, enhancing the model’s physical interpretability and accuracy.

Finally, we construct a residue-level graph where nodes represent residue features and edges encode the inter-residue geometries refined by the preceding CNN. Representation learning on this graph is performed using a graph transformer architecture ^[34]^. Each layer updates node embeddings according to **X**^(*l*+1)^ = LayerNorm(**X**^(*l*)^ + MultiHeadAttention(**X**^(*l*)^, E)), where **X**^(*l*)^ denotes the node features at layer *l* and E represents the edge features. This architecture computes multi-head attention scores that guide the aggregation of node information, facilitating efficient message passing across the entire graph structure.

### The atom-level graph learning

To explicitly model the influence of diverse sidechains on backbone conformation, we introduce an atom-level graph branch. We first construct graph representations for each of the 20 standard amino acids, with nodes as atoms and edges as covalent bonds. These individual amino acid graphs are then assembled into a full protein atom graph via peptide bond formation. This process is modeled as a dehydration reaction, wherein a water molecule (comprising two hydrogen and one oxygen atom) is eliminated upon the formation of each peptide bond (“–CO–NH–”) linking adjacent amino acids.

#### Algorithm 3 Atom Branch Graph Learning

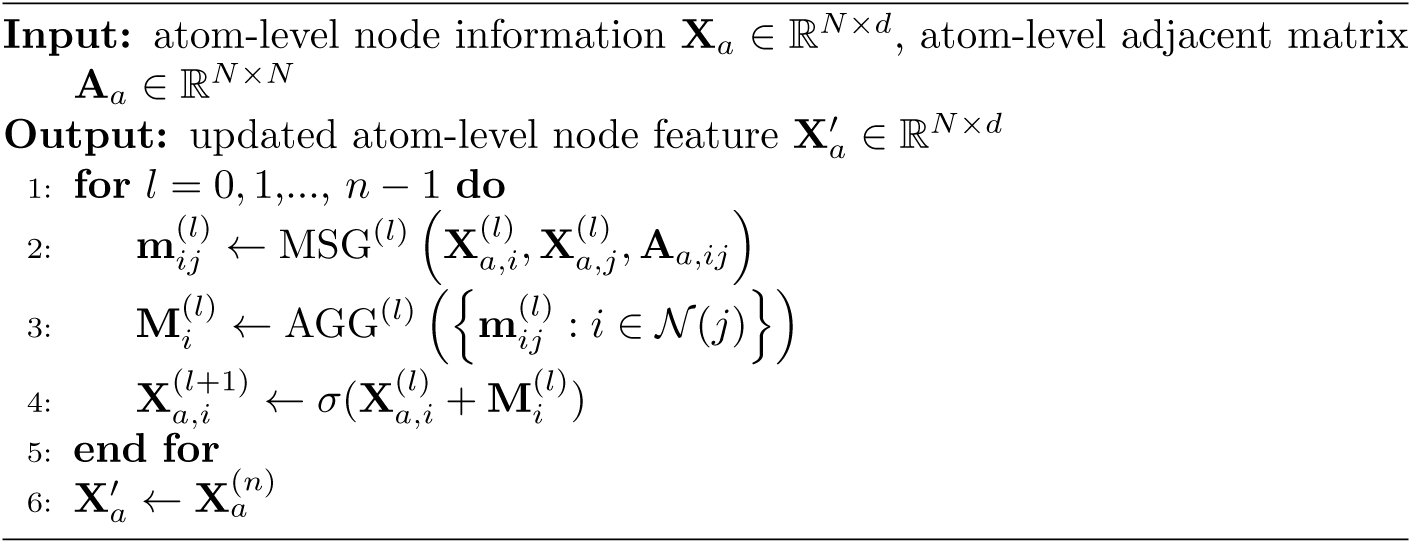

As shown in Algorithm 3, atom representations are then learned using a three-layer Graph Neural Network (GNN) block ^[35]^. Each layer updates the feature matrix as **X**^(*l*+1)^ = *GNN*(**A,X**^(*l*)^), where **X**^(*l*)^ is the feature matrix at *l*-th layer and **A** is the adjacency matrix. We incorporate residual connections between layers to enhance representational capacity. This architecture facilitates the propagation of sidechain information to backbone atoms, thereby enabling sidechain-dependent modeling of backbone conformation.

#### Algorithm 4 Residue-Atom Feature Fusion

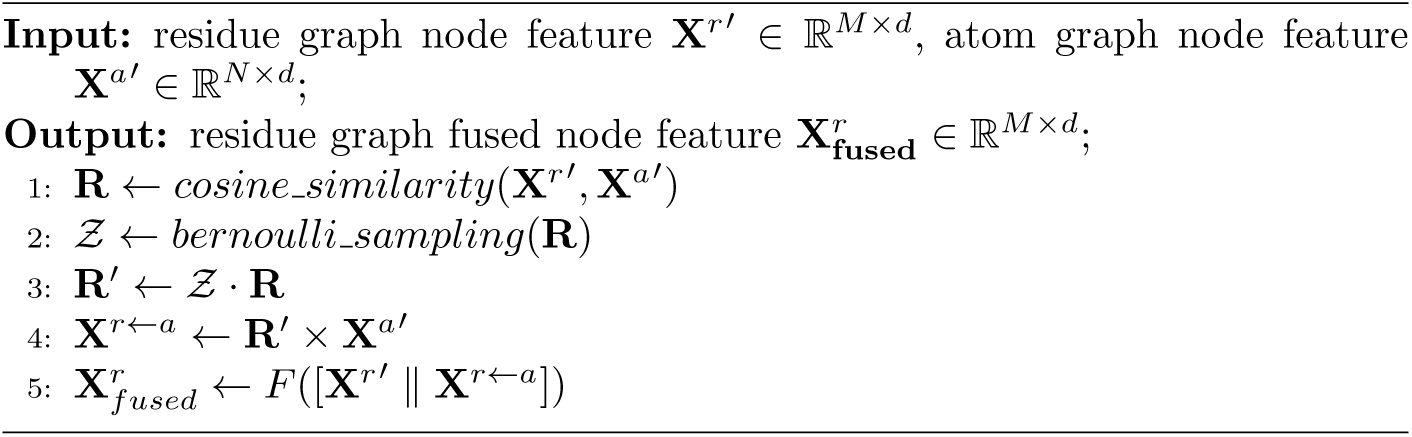

### The residue-atom graph feature fusion

We design a dual-level interaction learning process to learn weighting factors to fuse the residue-level and atom-level graphs. The residue features can be refined by mining atom-level features that affect the backbone conformation. Here, we use 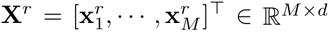 (*M* is the number of residues) to denote the residue feature matrix, and 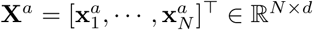 (*N* is the number of atoms) to denote the atom feature matrix.

In the dual-level interaction process (as shown in Algorithm 4), the residueatom relational matrix **R** ∈ R*^M^*^×*N*^ is first calculated, where the element in the *i*-th row and *j*-th column is formulated as 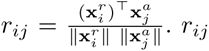. *r_ij_* represents the relationship between the *i*-th residue and *j*-th atom in the protein.

To massacre those unnecessary connections, we introduce a probabilistic sampling method to select critical atoms that influence protein residues. Concretely, given the residue-atom correlation **R**, we derive a set of latent random variables, denoted as Z, by learning the posterior distribution *p*(Z|**R**). Each element *z_ij_* ∈ Z follows a Bernoulli distribution *z_ij_* ∼ *B*(*p_ij_*), where *p_ij_* represents the probability that the *j*-th atom influences the folding of the *i*-th residue. The model then calculates an updated correlation matrix **R**^′^ via the element-wise product of the Bernoulli sample matrix Z and the original correlation matrix **R**.

Subsequently, the atom features **X***^a^* are projected into the space of residue features guided by the relationships in **R**^′^, yielding **X***^r^*^←*a*^ (as the mostly relevant atom for each residue). Finally, the residue features **X***^r^* are concatenated with the projected atom features **X***^r^*^←*a*^ to form an residue-atom interactive representation 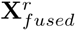, which is used for subsequent protein structure prediction.

### EGNN-based protein all-atom 3D-structure prediction

The 3D-structure predictor transforms the residue-atom fused features to all-atom coordinates through a two-stage inference (as shown in Algorithm 5). In the first stage, the SE(3)-EGNN ^[36]^ is used to project residue features into the rough 3D coordinates, i.e., the coordinates of backbone N, C*_α_* and C atoms for each residue.

Concretely, the initial backbone coordinates for the N, C*_α_* and C atoms (**bb coord^init^**) are computed by projecting the fused features through a multi-layer perceptron (MLP). A graph is then constructed where nodes represent residues, connected by edges to their top-k spatial neighbors and sequence-adjacent residues, as determined from these initial coordinates. This graph and the initial coordinates are processed by an SE(3)-EGNN to produce refined coordinates. The network computes self-attention scores between residue features, which are used to weight inter-residue messages. These messages are formulated using the direction vector **r***_ij_*, and are composed within an equivariant framework using spherical harmonics ^[48]^ and Clebsch-Gordan (CG) expansions ^[48]^. This architecture guarantees SE(3)-equivariance by construction, ensuring that rotations and translations of the input system produce consistently transformed outputs. Node features **h** are updated via message passing over node adjacent relations **A**. The final refined coordinates **bb coord^out^** are obtained by mapping these updated node features to coordinate displacements Δ**coord** and adding them to the initial coordinates **bb coord^init^**.

#### Algorithm 5 protein all-atom 3D structure predictor

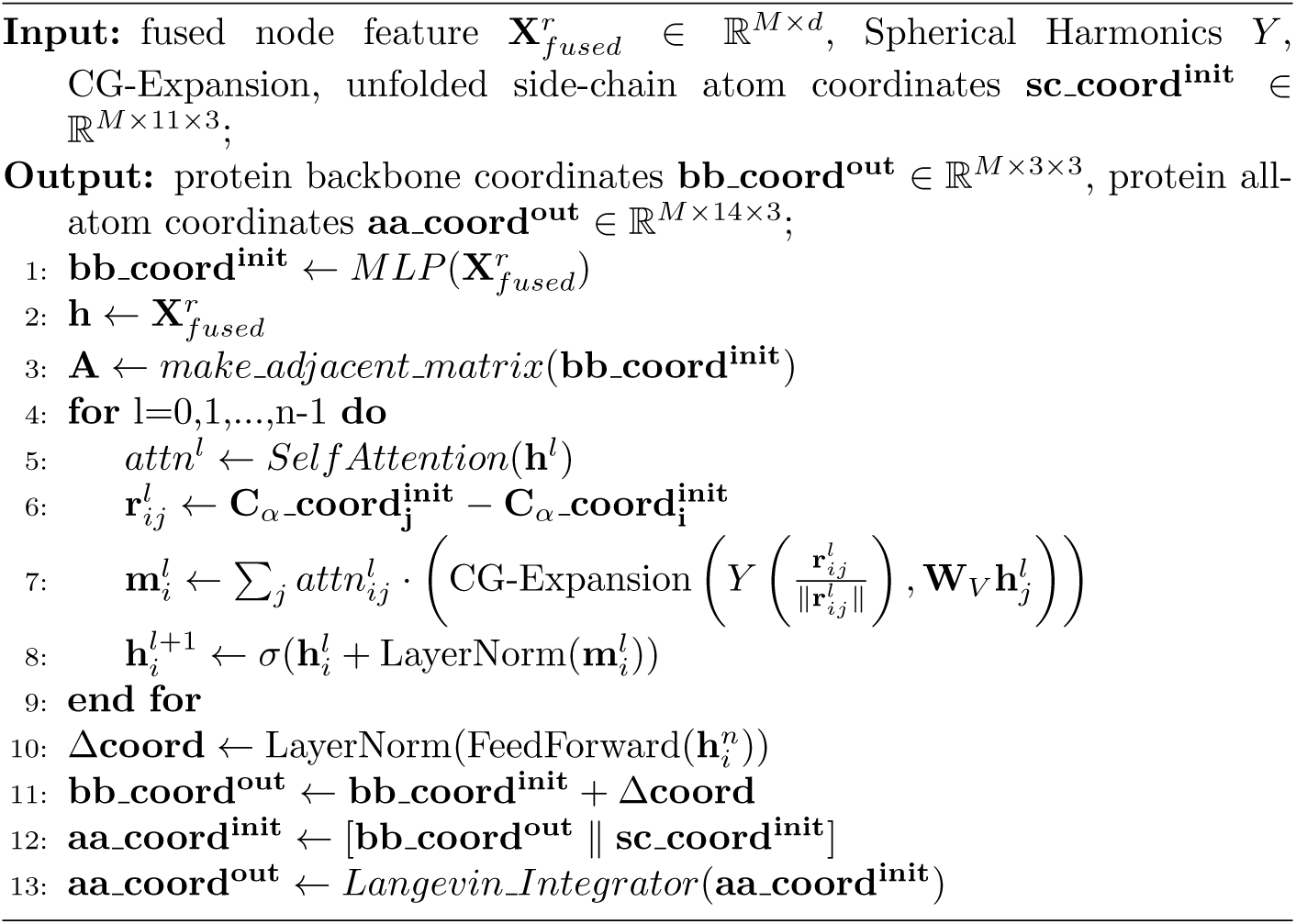

In the second stage, side-chain atoms are attached to the refined backbone. An all-atom structural optimization, inspired by molecular dynamics principles, is then applied to refine the side-chain conformations. This optimization mechanism operates with the backbone held fixed and comprises two key components: prior structural constraints that conduct optimal bond lengths, angles and dihedrals within the side chains, and a Langevin integrator ^[49]^ that introduces mild Brownian motion. The integrator prevents steric clashes by mitigating excessive atomic proximity that may arise during the optimized process of folding.

### Loss functions of sequence-geometry collaborative learning module

The training loss function of sequence-geometry collaborative learning (SCL) network, i.e. L*_bb_*, can be formalized as:

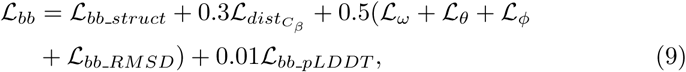

Where

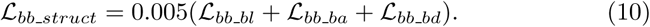

L*_bb___bl_* is the RMSE loss between bond length (N-C*_α_*, N-C, C*_α_*-C) computed from predicted coordinates of backbone structure and true bond length. For L*_bb___ba_* and L*_bb___bd_*, we compute the RMSE loss of bone angles (N-C*_α_*-C, C*_α_*-C-N, C-N-C*_α_*) and dihedrals (N-C*_α_*-C-N, C*_α_*-C-N-C*_α_*, C-N-C*_α_*-C), respectively.

L*_bb___RMSD_* is the coordinate RMSE loss between predicted and true backbone structure. L*_bb___pLDDT_* is the RMSE loss of LDDT score of C*_α_*.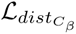, L*_ω_,* L*_θ_* and L*_ϕ_* are the cross-entropy loss for inter-residue distance and orientations.

In the all-atom training stage, sidechains are bonded to the backbone. Specifically, we set all these sidechain atom coordinates as optimizable parameters, and optimize them based on the following all-atom loss function:

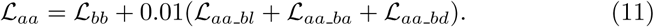

L*_aa___bl_,* L*_aa___ba_* and L*_aa___bd_* are the potential energy constrains (bond length, bond angle and dihedral) on sidechain and are also based on RMSE loss.

### Reporting summary

Further information on research design is available in the Nature Portfolio Reporting Summary linked to this article.

## 5 Data availability

The training structures of this study are acquired from the PDB (https://www.wwpdb.org/ftp/pdb-ftp-sites). For test benchmarks Orphan, Orphan25, CASP14, CASP15 and CASP16, they are freely available at https://figshare.com/s/57522d5f5aadc96f7e05.

## 6 Code availability

Source code for the TDFold model, trained weights, and inference script will be made available under an open-source license at https://github.com upon publication.

## Footnotes

∗ Different from the 3D structural template (homologous protein structure), 2D geometric template refers to pairwise distance and orientation matrices.

1 If additionally using MSA (i.e., *TDFold_MSA_*), the TM-score of TDFold can be further improved by 0.005-0.008 on the Orphan and Orphan25 datasets, but the inference time is about 10 times of the single sequence inference mode.

2 Although TDFoldMSA achieves an additional TM-score improvement of 0.02-0.03 on the CASP14, CASP15, and CASP16 datasets, the inference time is about 30 times of the single sequence inference mode.

3 Since DeepMind does not provide us with the model weights of AlphaFold3, we use the computation time of the AlphaFold3 server as the comparison result.

## References

[1] Jumper, J., Evans, R., Pritzel, A., et al: Highly accurate protein structure prediction with alphafold. Nature 596, 583–589 (2021)

[2] Baek, M., DiMaio, F., Anishchenko, I., et al: Accurate prediction of protein structures and interactions using a three-track neural network. Science. 117, 871–876 (2021)

[3] Zhang, J., Wang, Q., Barz, B., et al: Mufold: a new solution for protein 3d structure prediction. Proteins Struct. Funct. Bioinform. 78, 1137–1152 (2010)

[4] Hong, Y., Lee, J., Ko, J.: A-prot: Protein structure modeling using msa transformer. BMC bioinformatics. 23, 1–11 (2022)

[5] Zhang, C., Zheng, W., Mortuza, S., et al: Deepmsa: constructing deep multiple sequence alignment to improve contact prediction and fold-recognition for distant-homology proteins. Bioinformatics. 36, 2105–2112 (2020)

[6] Marks, D.S., Hopf, T.A., Sander, C.: Protein structure prediction from sequence variation. Nat. Biotechnol. 30, 1072 (2012)

[7] Ju, F., Zhu, J., Shao, B., et al: Copulanet: Learning residue co-evolution directly from multiple sequence alignment for protein structure prediction. Nat Commun. 12, 2535 (2021)

[8] Yang, J., Anishchenko, I., Park, H., et al: Improved protein structure prediction using predicted interresidue orientations. Proc. Natl. Acad. Sci. U. S. A. 117, 1496–1503 (2020)

[9] Xu, J., Mcpartlon, M., Li, J.: Improved protein structure prediction by deep learning irrespective of co-evolution information. Nature Machine Intelligence 3(7), 601–609 (2021)

[10] Abramson, J., Adler, J., Dunger, J., et al: Accurate structure prediction of biomolecular interactions with alphafold 3. Nature 630(8016), 493–500 (2024)

[11] Kryshtafovych, A., Schwede, T., Topf, M., et al: Critical assessment of methods of protein structure prediction (casp)—round xiv. Proteins: Structure, Function, and Bioinformatics 89(12), 1607–1617 (2021)

[12] Jeanmougin, F., Thompson, J., Gouy, M., et al: Multiple sequence alignment with clustal x. Trends in biochemical sciences 23(10), 403–405 (1998)

[13] Fiser, A.: Template-based protein structure modeling. Computational biology, 73–94 (2010)

[14] Suzek, B.E., Wang, Y., Huang, H., et al: Uniref clusters: a comprehensive and scalable alternative for improving sequence similarity searches. Bioinformatics. 31, 926–932 (2015)

[15] wwPDB Consortium.: Protein data bank: the single global archive for 3d macromolecular structure data. Nucleic Acids Res. 47, 520–528 (2018)

[16] Lin, Z., Akin, H., Rao, R., et al: Evolutionary-scale prediction of atomic-level protein structure with a language model. Science 379(6637), 1123–1130 (2023)

[17] Wang, W., Peng, Z., Yang, J.: Single-sequence protein structure prediction using supervised transformer protein language models. Nature Computational Science 2, 804–814 (2022)

[18] Chowdhury, R., Bouatta, N., Biswas, S., et al: Single-sequence protein structure prediction using a language model and deep learning. Nat Biotechnol 40, 1617–1623 (2022)

[19] Wu, R., Ding, F., Wang, R., et al: High-resolution de novo structure prediction from primary sequence. BioRxiv, 2022–07 (2022)

[20] Fang, X., Wang, F., Liu, L., et al: Helixfold-single: Msa-free protein structure prediction by using protein language model as an alternative. arXiv preprint arXiv:2207.13921 (2022)

[21] Vaswani, A., Shazeer, N., Parmar, N., et al: Attention is all you need. (2017). Paper presented at the 31st International Conference on Neural Information Processing Systems, Long Beach Convention Center, 4–9 December 2017

[22] Peng, J., Xu, J.: Raptorx: exploiting structure information for protein alignment by statistical inference. Proteins: Structure, Function, and Bioinformatics 79(S10), 161–171 (2011)

[23] Rombach, R., Blattmann, A., Lorenz, D., et al: High-resolution image synthesis with latent diffusion models. In: Proceedings of the IEEE/CVF Conference on Computer Vision and Pattern Recognition, pp. 10684–10695 (2022)

[24] Hu, E., Shen, Y., Wallis, P., et al: Lora: Low-rank adaptation of large language models. arXiv preprint arXiv:2106.09685 (2021)

[25] Ronneberger, O., Fischer, P., Brox, T.: U-net: Convolutional networks for biomedical image segmentation. In: Medical Image Computing and Computer-assisted intervention–MICCAI 2015: 18th International Conference, Munich, Germany, October 5-9, 2015, Proceedings, Part III 18, pp. 234–241 (2015). Springer

[26] Wang, X., Zhang, T., Liu, G., et al: Lightrosetta: High-efficient and accurate protein structure prediction using an ultra-lightweight deep graph model. bioRxiv, 2023–11 (2023)

[27] Kryshtafovych, A., Schwede, T., Topf, M., et al: Critical assessment of methods of protein structure prediction (casp)—round xv. Proteins: Structure, Function, and Bioinformatics 89(12), 1607–1617 (2023)

[28] Kryshtafovych, A., Schwede, T., Topf, M., et al: Critical assessment of methods of protein structure prediction (casp)—round xvi. Proteins: Structure, Function, and Bioinformatics 89(12), 1607–1617 (2025)

[29] Zhang, Y., Skolnick, J.: Scoring function for automated assessment of protein structure template quality. Proteins. 57, 702–710 (2004)

[30] Zemla, A.: Lga: a method for finding 3d similarities in protein structures. Nucleic acids research 31(13), 3370–3374 (2003)

[31] Mariani, V., Biasini, M., Barbato, A., et al: lddt: A local superposition-free score for comparing protein structures and models using distance difference tests. Bioinformatics 29, 2722–2728 (2013)

[32] Gutub, A.: Pixel indicator technique for rgb image steganography. Journal of emerging technologies in web intelligence 2(1), 56–64 (2010)

[33] Radford, A., Kim, J., Hallacy, C., et al: Learning transferable visual models from natural language supervision. In: International Conference on Machine Learning, pp. 8748–8763 (2021). PMLR

[34] Shi, Y., Huang, Z., Feng, S., et al: Masked label prediction: Unified message passing model for semi-supervised classification. (2021). Paper presented at the thirtieth International Joint Conference on Artificial Intelligence, virtual only, 19–26 August 2021

[35] Morris, C., Ritzert, M., Fey, M., et al: Weisfeiler and leman go neural: Higher-order graph neural networks (2019). Paper presented at the Conference of the Thirty-Third Conference on Artificial Intelligence, Hawaii, USA., 27 January – 1 February 2019

[36] Fuchs, F., Worrall, D., Fischer, V., et al: Se(3)-transformers: 3d rototranslation equivariant attention networks. (2020). Paper presented at the thirty-fourth Conference on Neural Information Processing Systems, virtual only, 6–12 December 2020

[37] Kullback, S., Leibler, R.A.: On information and sufficiency. Annals of Mathematical Statistics 22(1), 79–86 (1951)

[38] Lu, C., Zhou, Y., Bao, F., et al: Dpm-solver: A fast ode solver for diffusion probabilistic model sampling in around 10 steps. Advances in Neural Information Processing Systems 35, 5775–5787 (2022)

[39] Zhao, W., Bai, L., Rao, Y., et al: Unipc: A unified predictor-corrector framework for fast sampling of diffusion models. Advances in Neural Information Processing Systems 36, 49842–49869 (2023)

[40] Song, J., Meng, C., Ermon, S.: Denoising diffusion implicit models. arXiv preprint arXiv:2010.02502 (2020)

[41] Richardson, L., Allen, B., Baldi, G., et al: Mgnify: the microbiome sequence data analysis resource in 2023. Nucleic acids research 51(D1), 753–759 (2023)

[42] Altschul, S., Gish, W., Miller, W., Myers, E., Lipman, D.: Basic local alignment search tool. journal of molecular biology 215, 403–410, doi: 10.1016. S0022-2836 (05), 80360–2 (1990)

[43] Ho, J., Jain, A., Abbeel, P.: Denoising diffusion probabilistic models. Advances in neural information processing systems 33, 6840–6851 (2020)

[44] Chung, K.: Markov chains. Springer-Verlag, New York (1967)

[45] Norris, J.: Markov Chains vol. 2. Cambridge university press, London (1998)

[46] Zhai, J., Zhang, S., Chen, J., et al: Autoencoder and its various variants. In: 2018 IEEE International Conference on Systems, Man, and Cybernetics (SMC), pp. 415–419 (2018). IEEE

[47] Chen, S., Wong, S., Chen, L., et al: Extending context window of large language models via positional interpolation. arXiv preprint arXiv:2306.15595 (2023)

[48] Thomas, N., Smidt, T., Kearnes, S., et al: Tensor field networks: Rotation- and translation-equivariant neural networks for 3d point clouds. Advances in Neural Information Processing Systems (2018)

[49] Matthews, B.L.C.: Molecular Dynamics with Deterministic and Stochastic Numerical Methods. Springer (2015)

